# Underwater hyperspectral classification of deep sea corals exposed to 2-methylnaphthalene

**DOI:** 10.1101/150060

**Authors:** Paul Anton Letnes, Ingrid Myrnes Hansen, Lars Martin Sandvik Aas, Ingvar Eide, Ragnhild Pettersen, Luca Tassara, Justine Receveur, Stephane le Floch, Julien Guyomarch, Lionel Camus, Jenny Bytingsvik

## Abstract

Coral reefs around the world are under threat due to anthropogenic impacts on the environment. It is therefore important to develop methods to monitor the status of the reefs and detect changes in the health condition of the corals at an early stage before severe damage occur. In this work, we evaluate underwater hyperspectral imaging as a method to detect changes in health status of both orange and white color morphs of the coral species *Lophelia pertusa*.

Differing health status was achieved by exposing 66 coral samples to the toxic compound 2-methylnaphthalene in concentrations of 0 mg L^*−*1^ to 3.5 mg L^*−*1^. A machine learning model was utilized to classify corals according to lethal concentration (LC) levels LC5 (5 % mortality) and LC25 (25 % mortality), solely based on their reflectance spectra. All coral samples were classified to correct concentration group. This is a first step towards developing a remote sensing technique able to assess environmental impact on deep-water coral habitats over larger areas.

## Introduction

Coral reefs are the foundation of many marine ecosystems. Corals are found across the world’s ocean, in both shallow tropical and subtropical waters and in deep water. Deep-water corals thrive in cold, dark water at depths of up to 6000 m [1]. They have a cosmopolitan distribution, being particularly abundant in the North East Atlantic, and are for instance found off the coast of Norway and deep underwater in the Mediterranean Sea. In contrast to tropical corals, they are azooxanthellate, meaning that they do not have symbiotic life forms with dinoflagellates and hence do not require direct access to sun light [2]. *Lophelia pertusa* (Linneaeus 1758) is one of the most abundant reef forming scleractinian deep-water corals in cold and temperate regions, with main occurrences at depths ranging from 200 m to 1000 m [3, 4]. The species is found as two color morphs (orange and white) due to different pigment composition causing phenotype specific optical fingerprints [5, 6]. The reef framework offers structural habitat for a variety of benthic species [2], including gorgonian corals, sponges, squat lobsters (*Munida sarsi*) and rosefish (*Sebastes viviparoius*) as well as economically important fish species such as atlantic cod (*Gadus morhua*), saithe (*Pollachius virens*), and cusk (*Brosme brosme*) [3, 7–9].

The coral reefs are in danger, primarily due to climate change and increased CO_2_ levels [10], leading to ocean acidification. The increased CO_2_ levels can affect the ability of the coral to form its aragonite skeleton [11, 12]. The combined pressures can result in coral bleaching, slower growth [13] and reproduction rates, and degraded reef structures [14]. Other chemical and physical stressors may also damage corals. The oil industry is moving northwards and many of the drilling areas coincide with habitats listed by the Convention for the Protection of the Marine Environment of the North-East Atlantic (OSPAR) as being potentially rare or declining, including the cold-water coral reefs and sponge grounds. Corals and sponges are slow growing [15], and their habitats are therefore regarded as vulnerable to the smothering effects from drill cuttings [16]. Coral colonies affected by the Deepwater Horizon oil spill have shown signs of stress and mortality [17, 18]. Mechanical damages from trawling vessels are also considered a threat to the reefs [3, 15].

The Norwegian environmental regulatory authorities have imposed requirements for detailed habitat and organism mapping prior to exploratory drilling, as well as post-drilling surveying to map the distribution of deposited drill cuttings and the extent of possible biological impacts. Further, mapping of sensitive species and habitats to accidental oil pollution is an essential part of contingency plans, and species distribution maps are a crucial tool to assist responders during an incident, especially when the industrial activities are near shore.

Hyperspectral imaging collects and processes incoming light across a range of the optical spectrum. In contrast to ordinary photographic cameras (or similarly, the human eye) which record only three color bands (red, green, and blue), a hyperspectral camera produces a full spectrum of all available wavebands in each pixel in an image. Objects will reflect and absorb light to a varying degree at different wavelengths depending on their color and pigmentation providing a spectral signature of that object. Hyperspectral imaging generates large amounts of data which require sophisticated data analysis and machine learning methods [19]. Multivariate data analysis and machine learning have been used successfully in several marine environmental studies for the interpretation of large data sets, for example in integrated environmental monitoring [20] and for the analyses of photos to assess the potential impact of water-based drill cuttings on deep-water rhodolith-forming calcareous algae [21].

Hyperspectral imaging is widely used as an in-situ and non-invasive sensor for species mapping and detection of ecosystem health status [22]. Hyperspectral imagers are typically operated from satellite or aeroplane [23], for instance, shallow water corals are monitored by remote sensing from satellites [24]. The Underwater Hyperspectral Imager (UHI) used in the present work represents a recent system for identification, mapping and monitoring of objects of interest (OOI) at the seabed [25, 26]. However, underwater spectral measurements have been used to measure changes in coral appearance using point measurements [27], multispectral camera [28] and hyperspectral camera [29], all in the visible part of the electromagnetic spectrum, as both ultra violet and infrared radiation is attenuated in water.

The purpose of the present work was to evaluate the use of UHI and multivariate data analysis to detect changes in health condition of the coral species *L. pertusa*. Corals were exposed to 2-methylnaphthalene in laboratory experiments in order to provide corals with health condition varying from unaffected to dead. Hyperspectral images of exposed and control corals were then recorded after a recovery period. Finally, classification of these images using machine learning shows in a visual way which spatial areas are affected by exposure to toxic compounds.

## Materials and methods

In order to evaluate the use of underwater hyperspectral imaging as a *L. pertusa* health detection tool, corals needed to be sampled and different health conditions established prior to acquisition of hyperspectral images. Hence, the experimental work presented in this study consists of the following activities: collecting and rearing of coral samples, exposure to the toxicant 2-methylnaphthalene, monitoring the corals to determine polyp mortality, and imaging them using UHI. An overview of the timeline is given in Table 1.

**Table 1.**
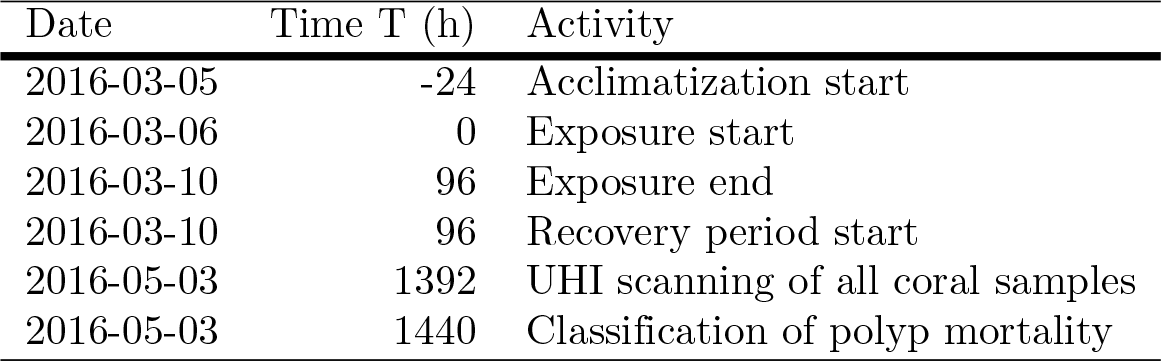
Timeline of experimental activities.

### Sampling and rearing of corals

Samples of *L. pertusa* were collected at Stokkbergneset in Trondheimsfjorden (Norway, 63.47° N, 9.91° E) on September 1st 2015 in collaboration with the Norwegian University of Science and Technology (NTNU) onboard R/V Gunnerus. The site is characterized by a steep rock wall with *L. pertusa* occurring in colonies from 100 m to 500 m depth [30].

The coral samples were collected from four different colonies at depths between 208 m to 235 m. Both white and orange color morphs were sampled. A Sperre Subfighter 7500 remotely operated vehicle. (ROV) (“Minerva”) with an attached fish net was used to conduct the sampling. Precaution was taken in order to avoid any damages on other parts of the reefs. This is described in more detail in S5 Fig.

*L. pertusa* has a wide geographical distribution throughout the world, and the density is particularly high along the coast of Norway. The species is not defined as threatened or protected in Norway, but due to the reported decline the last 10 years, they are defined as nearly threatened according to the terms used in the Norwegian Red List for Species. The list is prepared by the Norwegian Biodiversity Information Centre in accordance to the International Union for Conservation of Nature (IUCN). As the sampling was conducted outside of national parks or other type of protected area, no specific permissions were required.

The corals were transported to the Akvaplan-niva Research and Innovation Center Kraknes (RISK) (Tromsø, Norway) by car in a water tank where temperature, oxygen levels and oxygen saturation were frequently monitored. The temperature range and oxygen saturation during transport was 5.1 ^*◦*^C to 9.8 ^*◦*^C and 98 % to 113 %, respectively. The pieces of *L. pertusa* were divided into smaller samples and placed in silicon tubes attached to steel grids (20 cm × 40 cm) in 500 L tanks with a continuous flow (500 L h^*−*1^) of filtered (60 µm filter) and UV-treated bottom sea water (63 m depth) from the fjord Sandøysundet, adjacent to the research facility. One sample refers to a coral branch with 3 to 9 polyps, as shown in Fig 2. The temperature, oxygen saturation, and oxygen concentration were monitored on a daily basis during rearing, and the corals were fed three times per week with *Calanus* sp. nauplii from Planktonic AS. The corals were cleaned from sediments once per week.

### Exposure setup and procedure

2-methylnaphthalene is classified as a low molecular weight poly aromatic hydrocarbon (PAH) with high water solubility. The compound was chosen because it was expected to be sufficiently water soluble and toxic to produce corals with varying health status. In addition, PAHs are abundant constituents in crude oil and are also found in water from oil wells [31].

The set-up comprised five treatment groups: one control group (C0) and four exposed groups (C1, C2, C3 and C4). Each treatment group consisted of four replicates (R1, R2, R3 and R4). Each replicate consisted of one orange and two white pigmented morphs, giving a total of 20 orange and 40 white coral samples. The corals were assessed to be in good condition when sampled. The samples were randomly distributed in order to represent all four coral colonies at Stokkbergneset.

Each sample comprised a coral branch with 3 to 9 polyps. Briefly, after 24 h acclimatization, *L. pertusa* were used in a 96 h acute toxicity testing to 2-methylnaphthalene (see Table 1 for a timeline). During acclimatization and exposure, each coral replicate were kept in a square glass aquarium containing 1.5 L sea water. The animals were not fed during the toxicity testing.

The acute toxicity tests to 2-methylnaphthalene involve passive dosing. Based on the method described by Butler et al. [32], a passive dosage system using silicone O-rings has been developed. This was done to obtain a stable concentration of 2-methylnaphthalene as close to the nominal concentration as possible. The O-rings were cleaned before they were loaded with 2-methylnaphthalene with target concentrations in seawater of 0 mg L^*−*1^, 1.03 mg L^*−*1^, 2.27 mg L^*−*1^, 5.00 mg L^*−*1^, and 8 mg L^*−*1^ to 10 mg L^*−*1^ (saturation) for C0 to C4, respectively. After O-rings were loaded, they were set to equilibrate with seawater in 20 L bottles for 24 h, one bottle per treatment group. The final stock solution was pumped into the exposure chambers continuously using a peristaltic pump with the inlet placed in the bottom of the exposure beakers as illustrated in Fig 1. Overflow ensured that the overhead between water surface and glass lids covering the exposure beakers always was at a minimum. Using O-rings combined with a pump is beneficial as it reduces the interference and stress on the corals during the exposure period. Based on a combination of the coral oxygen needs, chemistry and kinetics, tubing capacity, pump capacity and practical reasons, a flow rate of approximately 160 mL h^*−*1^ was selected. The exchange rate of exposure solvent in the exposure beakers was about 2.5 times per 24 h. In addition to logging of temperature every 30 min throughout the 96 h exposure (Tidbit Temperature Data Logger V2, Onset, Massachusetts), the temperature and oxygen saturation was measured daily throughout the experiment (Oxyguard). The chambers were covered with a glass lid to reduce evaporation of 2-methylnaphthalene. All stock solution bottles and treatment group replicates were placed on a magnet stirrer throughout the whole experiment to keep the exposure media homogeneous. The exposure was conducted in a room holding 4 ± 1 ^*◦*^C. After the toxicity test was ended, all coral samples were kept for approximately 12 h in individual tanks with pure seawater to provide an initial period of depuration of 2-methylnaphthalene before they were transferred to a single recovery tank of the same type and with similar facilities as they were kept in prior to the toxicity test.

**Fig 1.** Setup for 2-methylnaphthalene exposure. A peristaltic pump circulates stock solution to the exposure chamber.

In addition to the 60 samples in the five treatment groups, six coral samples (two orange and four white) referred to as reference alive were also included in the set-up. The reference alive corals originated from the same coral colonies as the samples used in the toxicity test. The reference samples were kept in the rearing tanks with pure seawater while the C0–C4 corals went through the toxicity experiment. Reference alive corals were kept in the same tank as C0–C4 during the recovery period. Hence, reference alive corals were exposed to minimal handling compared to C0–C4 before they were imaged with the UHI. The reference alive group was included as an additional control group in case of accidental contamination of the experimental control group (C0) or in case handling itself affected the corals and their spectral properties.

### Polyp mortality

Prior to exposure to 2-methylnaphthalene (at time –24 h, see Table 1 for a timeline), the number of alive polyps on each coral sample was counted. Mortality was assessed after recovery and UHI scans (at time 1440 h). This approach was chosen because it was not possible to determine whether polyps were alive or dead immediately after end of exposure. By keeping the coral samples in a recovery tank over weeks, the samples and individual polyps could be monitored visually for the presence of soft tissue on skeleton and polyps. A polyp was classified as dead when soft tissue was no longer present within the polyp’s calyx. Illustration of live and dead polyps are given in Figure 2. The fraction of dead polyps (i.e., the number of dead polyps divided by the number of alive polyps before toxicant exposure) is presented in results and statistics as polyp mortality.

**Fig 2.** Example images of living and dead corals.

Example images of alive and dead polyps. Upper row – pictures from time 0 h: orange and white coral samples with healthy polyps. Lower row – pictures from time 1392 h: orange and white coral samples with dead polyps (red ellipse indicates dead polyps). The coral sample in the lower left picture was orange prior to exposure, but exposure to 2-methylnaphthalene lead to loss of pigmented tissue and has rendered it predominantly white. The lower right picture exhibits a white coral sample with dead polyps.

### Chemical analysis of exposure concentration

#### Preparation of water samples

10 mL water samples were collected from the exposure beakers in Tromsø, Norway. Samples were acidified to pH 1 using hydrogen chloride (HCl), added methanolic solution (1.8 mL) of perdeuterated naphthalene (Nd8) at the concentration of 100 µg mL^*−*1^ and frozen at –20 ^*◦*^C. The final concentration of Nd8 in water samples was 15.3 µg mL^*−*1^. The water samples were then sent to Brest, France in a cool box for analysis.

#### Extraction of water samples and conditions of analysis

The water samples were thawed overnight in the fridge, and then extracted with 2 mL of pentane. A calibration curve was established for 2-methylnaphthalene concentrations in the range 0 mg L^*−*1^ to 10 mg L^*−*1^ with a fixed concentration of Nd8.

Samples were analyzed by Gas Chromatography coupled to Mass Spectrometry (GC-MS). The GC was an HP 7890 series II (Hewlett-Packard. Palo Alto. CA. USA) equipped with a Multi Mode Injector (MMI) used in the pulsed splitless mode (Pulse Splitless time: 1 min. Pulse Pressure: 103 kPa). The injector temperature was maintained at 300 ^*◦*^C and 1 µL of the extract was injected. The interface temperature was 300 ^*◦*^C. The GC temperature gradient was from 70 ^*◦*^C (0 min) to 250 ^*◦*^C (0 min) at 15 ^*◦*^C min^*−*1^. The carrier gas was Helium at a constant flow of 1 mL min^*−*1^. The capillary column used was a HP 5-ms (Hewlett–Packard, Palo Alto. CA. USA, 30 m in length, 0.25 mm internal diameter, and a film thickness of 0.25 µm). The GC was coupled to a HP 5975 Mass Selective Detector (MSD) used in the Electronic Impact mode (electronic impact energy 70 eV, system temperatures: 230 ^*◦*^C (source) and 150 ^*◦*^C (quadrupole)). 2-methylnaphthalene quantifications were done using Single Ion Monitoring mode with respectively the molecular ion of each compound at a minimum rate of 2 s^*−*1^ (mass/charge ratio of 142 for 2-methylnaphthalene and 136 for Nd8). 2-methylnaphthalene was quantified relatively to the perdeuterated Nd8 introduced at the beginning of the sample preparation procedure using calibration curves.

### Underwater hyperspectral imaging

The underwater hyperspectral imager (UHI) is a line camera (often referred to as a “push broom” sensor), consisting of a narrow slit, a spectrograph, and a monochrome 2D camera, mounted in a waterproof housing made of aluminum with a fused silica window. Communication with the camera is achieved through a sub-sea cable.

#### Image acquisition

For capturing images of an area, the UHI is installed on a moving platform, where it captures frames perpendicular to its direction of motion. The UHI was configured to acquire data in the wavelength range 381 nm to 846 nm.

Reflectance is a physical property of an object that can be attributed to material properties. The measured reflectance depends on several parameters such as incident and observation angle, and hence the surface structure of the material in interest. In remote sensing and imaging it is common to assume diffuse reflectance from the targets. This is an assumption about the smoothness of the surface of materials which is surprisingly often valid in nature.

In water, the largest obstacle for achieving accurate reflectance measurements is the optical wavelength dependent attenuation of the water itself. This property alters the apparent color of an object if it is viewed through water at varying depths (optical path length). Water from different sources in time and space can have non-identical constituents and hence a different attenuation coefficient.

To compensate for the optical attenuation of water, a Spectralon diffuse reflectance standard (Labsphere inc) with dimensions 20 cm *×* 20 cm *×* 3 cm was imaged alongside the coral samples. By comparing the measured spectra of the Spectralon, *I*_spec_(*x, λ*), with its calibrated reflectance spectrum, *R*_spec_(*λ*), a conversion factor, *A*(*x, λ*), was found for every spatial pixel (at position *x*) covered by the Spectralon.

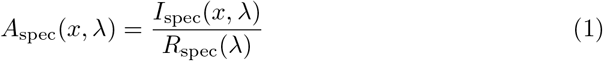

Due to the small size of the reference target a larger PVC plate, roughened with a 500 grit size sandpaper, was used in addition. The reflectance spectrum of the reference target was used to calculate the reflectance spectrum of the PVC plate. The reflectance of the PVC, *R*_PVC_(*λ*), was then calculated by applying the conversion factor, *A*_spec_(*x, λ*), to the spatial pixels in the UHI slit covering the Spectralon, when imaging the PVC. Since Spectralon was 3 cm thick, the reflectance of the PVC plate was calculated for 3 cm as (*N*_spec_ is the number of Spectralon pixels):

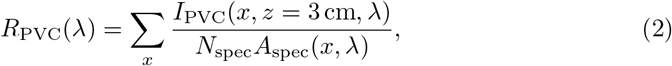

In order to account for the optical path length of samples of various heights the PVC plate was inclined such that all spatial pixels could be compared to a known reflectance at its own height.

The conversion factors could then be found for all spatial pixels in the slit at any altitude,

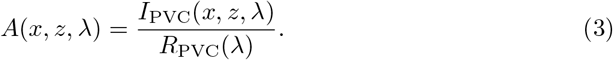

The height of the coral samples was measured to be approximately 7 cm using a ruler. The reflectance of the coral samples could thus be obtained using

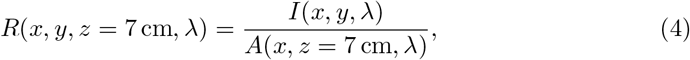

where *I*(*x, y, λ*) is the recorded intensity, i.e., the hyperspectral image.

The coral samples and reflectance references were placed in a tank of dimensions 2.0 m 1.0 m 1.6 m, filled with sea water. The UHI and lighting was mounted on a linear motorized rail; see Fig 3 and S6 Fig. The vertical distance between the spectrograph and coral samples was 0.35 m. Two halogen lamps (Osram Decostar 51 TITAN 50 W 12 V 60° GU5.3) with constant power supply were the only light sources during the measurements. Halogen lights were used because of their relative good uniformity across the wavelengths of light for which water is transparent. The whole setup was surrounded by a black tent to avoid light pollution. The UHI was moved along the rail with a constant speed of 0.13 m s^*−*1^ during imaging. By comparison with focal sheets placed in the bottom of the tank, the spatial resolution of the recorded images was estimated to be approximately 2.5 mm in the center of the slit. The diffraction limited spectral resolution was approximately 5 nm, while the spectra were sampled approximately every 0.5 nm.

**Fig 3.** Experimental setup for UHI image acquisition.

Samples of *L. pertusa* were set up on the bottom of a tank and imaged using an underwater hyperspectral imager (UHI). The UHI was attached to a linear scanning mechanism and operated in a “push-broom” fashion. The setup also includes a Spectralon reference plate and an inclined reference plate to account for changes in the spectrum due to the presence of water.

### Pixel extraction

The reflectance spectra from each coral sample was extracted by manually selecting image pixels; a total of 103 to 718 pixels were selected depending on the size of the coral. Pixels were selected both from the branches and the polyps of the coral. The reflectance spectrum was labelled with the sample ID and stored as an entry in a database. Each entry contained information about the measured reflectance for each pixel at each wavelength from 381 nm to 846 nm, and the pixel position in the original UHI image (corresponding to the position in the tank).

### Data used in spectral classification

Spectra were recorded for the 60 coral samples representing the C0 to C4 corals used in the exposure study, as well as for the 6 reference alive corals. Due to poor transmission of ultra violet and near infrared wavelengths even in extremely clean water [33], poor signal quality was experienced in both ends of the spectrum. Therefore, only data from the spectral range 400 nm to 750 nm are included in this study. The spectra recorded are placed in a matrix **X**, where each column corresponds to one wavelength, and each row corresponds to one observation, i.e., a pixel in the hyperspectral image.

The matrix **Y**_*h*_, used for Projection to Latent Structures (PLS) [34] in the *preprocessing* stage only, consists of the following columns: 2-methylnaphthalene concentration and polyp mortality. Each row of the matrix **Y**_*h*_ corresponds to the same row in **X**, i.e., spectra associated with the same coral sample.

The vector **Y**_*c*_, used for *classification*, consists of one categorical variable, namely, the exposure category of the organism which the spectrum came from. The exposure category is determined by the LC5 and LC25 concentration (lethal concentration for 5 % and 25 % of polyps) of 2-methylnaphthalene: the *non-or low* exposure category was exposed to a concentration *C* ≤ LC5; the *intermediate* exposure category was exposed with LC5 *< C* ≤ LC25; and the *high* exposure category was exposed with *C >* LC25.

### Classification algorithm

A three-stage machine learning model was applied to classify the exposure level of the coral samples. A flow chart for the data analysis process is given in Fig 4, showing the steps in the data analysis. In all cases, separate models were trained on spectra from white and orange corals, as the pigmentation of the coral can easily be determined using a simple linear classifier

**Fig 4.** Data analysis flowchart.

The flowchart shows the full data analysis pipeline including all steps of the classification model.

The stages consist of standardization, PLS and transformation, followed by a *ν*-Support Vector Machine (*ν*SVM) classification algorithm [35–37]. Note that linear classification algorithms were found to be unsuitable to the problem at hand, due to the nonlinear separation of points in the PLS latent variable space. The scikit-learn software package was used for both of the PLS and SVM algorithms [35].

#### Standardization of spectra

Before attempting classification, the model inputs were standardized:

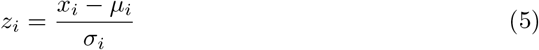

where *z_i_* is the scaled value, *x_i_* is the original value, and *µ_i_* and *σ_i_* is the mean and standard deviation of feature *i* (i.e., the spectrum intensity at wavelength *i*), respectively. Standardization implies that all input variables to the model have zero mean and unit variance, thus avoiding influence of, e.g., light intensity variations caused by placement of light sources, unrelated to the reflectance of the imaged object. Each feature corresponds to a wavelength in the spectrum, and a column of the input data matrix **X**:

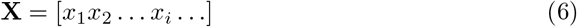

where the *x_i_* are column vectors containing all measurement values. Similarly, standardization was also performed on **Y**_*h*_ (2-methylnaphthalene concentration and polyp mortality).

#### Dimensionality reduction

Following standardization of inputs, a PLS model relating spectra (**X**) to 2-methylnaphthalene concentration and polyp mortality (**Y**_*h*_) is constructed:

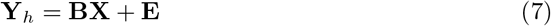

where **E** is the error term. In the process of constructing the regression coefficient matrix, **B**, the input matrix **X** is transformed to a lower-dimensional (latent) subspace; hence, the name PLS. This is done in a way such that the covariance between the **X** and **Y**_*h*_ matrices is maximized. Furthermore, PLS selects variables that extract the maximum relevant information possible. These *latent variables* are simply weighted sums of the original variables (e.g., the recorded spectrum intensity at each wavelength). One can think of the colors red, green, and blue as three latent variables used by our eyes to interpret the optical spectrum. The vector of the weights are referred to as *loading vectors* and reveals which underlying variables (in our case, wavelengths) that have the most explanatory power. Note that the loadings are ordered such that the first PLS component contains the most explanatory power, and subsequent component contain less and less explanatory power between the variables. In our case, a dimensionality reduction from 619 dimensions (wavelengths) to 10 dimensions (i.e., 10 latent variables) was chosen. The number of latent variables was selected using 5-fold stratified cross-validation [38]; however, we note that the results were not sensitive to the exact number of latent variables, and that classification performed well for 5 to 20 latent variables.

Subsequently, classification is improved both in quality and speed, as noise is removed and the dimensionality of input data is reduced significantly. Also, separation of spectra is improved as the PLS algorithm maximizes covariance between spectra and 2-methylnaphthalene concentration and polyp mortality.

#### Classification of spectra and samples

The input to the classification stage is, as described above, spectra (**X**) transformed into the 10-dimensional PLS latent variable space, as well as 2-methylnaphthalene concentration and polyp mortality (**Y**_*c*_). The *ν*SVM model constructed uses the parameter *ν* = 0.1 and a radial basis function kernel [36], as the spectra are not linearly separable in the latent variable space.

After classification of individual spectra, we classify each coral sample to the class with the highest number of single spectra (the majority vote algorithm). This lends significant robustness to incorrect classification, as many individual spectra can be classified incorrectly without affecting overall area coverage.

We note that model training and testing takes on the order of one minute on a desktop computer, and the use of the model to classify new data require only fast and well optimized computational operations, e.g., matrix multiplication. As such, the computational requirements are light, and should not pose any problems for practical use.

## Results and discussion

### Chemical analysis of 2-methylnaphthalene concentration

Chemical analysis of water samples from the stock bottles confirm that the desired concentrations of 2-methylnaphthalene, by using loaded silicon O-rings, were obtained. The average 2-methylnaphthalene concentration in the water samples from the stock bottles sampled at time 0 h, 24 h, 48 h and 72 h were 125 %, 115 %, 70 % and 98 % of nominal concentrations for C1 to C4, respectively (see Table 2 and S1 Table). The 2-methylnaphthalene levels in the C0 stock bottles samples at the same time points were all below 0.033 mg L^*−*1^ (which was the limit of detection), and hence, regarded as acceptable. The 2-methylnaphthalene concentration in the water samples from the exposure beakers sampled from T0 to T72 were somewhat lower than in the stock bottles with C1 to C4 calculated to be 43 % (replicates ranging from 38 % to 51 %),47 % (replicates ranging from 40 % to 60 %), 27 % (replicates ranging from 2 % to 47 %) and 32 % (replicates ranging from 26 % to 42 %) of nominal concentrations, respectively; see Table 2 and S1 Table. However, one of the four C3 replicates (R2) deviated from the others. The replicate C3 R2 contained significantly lower 2-methylnaphthalene concentrations than the other replicates with only 2 % of nominal concentration achieved, while the remaining three replicates ranged from 39 % to 47 % of the nominal value (see S1 Table). The deviation of C3 R2 was probably due to a faulty pump channel. The levels of 2-methylnaphthalene in the control (C0) beakers were all below 0.07 mg L^*−*1^, and hence, regarded as acceptable. The lower concentration of 2-methylnaphthalene in the exposure beakers compared to the stock bottles is likely explained by evaporation of 2-methylnaphthalene from water masses to air as the systems were not completely closed. As stock solutions were pumped through the tubes connecting the stock bottle and the exposure beakers 24 h prior to exposure start, and as teflon coated tubes were used, we believe that there was no loss of 2-methylnaphthalene during the 96 h exposure period due to 2-methylnaphthalene sticking to the tube walls. It is unlikely that bioaccumulation of 2-methylnaphthalene in coral tissue was of significance as the biomass of corals in each replicate beaker was low (ranging from 9 g to 21 g coral, skeleton and soft tissue, in 1.5 L sea water). In all beakers, the concentration of 2-methylnaphthalene was lower at T0 than measured at later time points. This initial increase is explained by non-contaminated water masses being replaced by contaminated stock solution during the first hours (flow of approximately 160 mL h^*−*1^) of the exposure period. Besides this, the concentration of 2-methylnaphthalene in the exposure beakers correlates with the concentrations in the stock solutions (Pearsons’ correlation *R*^2^ = 0.61, *p* ≤ 0.001).

**Table 2.** Concentration of 2-methylnaphthalene (mg L^*−*1^.

| Exposure group |  | C0 | C1 | C2 | C3 | C4 |
| --- | --- | --- | --- | --- | --- | --- |
| Nominal concentration (mg L<sup>-1</sup>) |  | 0.0 | 1.0 | 2.3 | 5.0 | 8.0 |
| Stock solution | avg | 0.0 | 1.3 | 2.6 | 3.5 | 7.8 |
|  | std dev | 0.0 | 0.2 | 0.5 | 1.3 | 1.2 |
|  | min | 0.0 | 1.0 | 1.9 | 1.8 | 6.1 |
|  | max | 0.0 | 1.5 | 3.2 | 5.0 | 10.2 |
| Beakers 1–4 (avg) | avg | 0.0 | 0.4 | 1.1 | 1.3 | 2.6 |
|  | std dev | 0.0 | 0.2 | 0.6 | 1.7 | 1.4 |
|  | min | 0.0 | 0.0 | 0.0 | 0.0 | 0.0 |
|  | max | 0.1 | 0.8 | 2.0 | 8.9 | 4.9 |
| Beaker 1 | avg | 0.0 | 0.4 | 1.4 | 1.4 | 2.6 |
|  | std dev | 0.0 | 0.3 | 0.7 | 0.8 | 1.2 |
|  | min | 0.0 | 0.0 | 0.4 | 0.4 | 1.0 |
|  | max | 0.0 | 0.6 | 2.0 | 2.3 | 3.9 |
| Beaker 2 | avg | 0.0 | 0.4 | 1.0 | 0.1 | 2.4 |
|  | std dev | 0.0 | 0.2 | 0.4 | 0.2 | 1.6 |
|  | min | 0.0 | 0.1 | 0.3 | 0.0 | 0.0 |
|  | max | 0.0 | 0.7 | 1.3 | 0.5 | 4.2 |
| Beaker 3 | avg | 0.0 | 0.4 | 1.0 | 2.4 | 3.4 |
|  | std dev | 0.0 | 0.2 | 0.6 | 2.8 | 1.5 |
|  | min | 0.0 | 0.1 | 0.0 | 0.4 | 1.0 |
|  | max | 0.0 | 0.7 | 1.5 | 8.9 | 4.9 |
| Beaker 4 | avg | 0.0 | 0.5 | 0.9 | 1.4 | 2.0 |
|  | std dev | 0.0 | 0.3 | 0.6 | 0.9 | 1.4 |
|  | min | 0.0 | 0.1 | 0.0 | 0.4 | 0.0 |
|  | max | 0.1 | 0.8 | 1.4 | 2.7 | 3.5 |

### Polyp mortality

The fraction of dead polyps at the end of the experiment correlated with the 2-methylnaphthalene exposure doses, and based on the sigmoidal dose-response curve the LC5 and LC25 values were determined (Fig 5). Each point in the figure represent one sample (with 3 to 9 polyps), with groups of 3 samples sharing the same 2-methylnaphthalene concentration, as they were placed in the same exposure beaker. Concentrations of 2-methylnaphthalene are based on levels in water samples collected during the exposure period, while the polyp mortality was assessed approximately 8 weeks later. This approach was necessary because it was challenging, if not impossible, to determine whether polyps were alive or dead shortly after exposure was ended. The coral samples could have both live and dead polyps at the time they were assessed. Whether mortality of one single polyp has any impact on neighboring polyps are unknown as there to our knowledge is highly limited knowledge about communication and signaling between polyps of *L. pertusa*. However, it has been shown that there are no nervous communication between polyps in *L. pertusa* colonies [39]. Although no firm conclusion can be given, we believe that there has been no influence of intercommunication between polyps affecting our mortality data.

**Fig 5.** Dose-response curve for 2-methylnaphthalene exposure and polyp mortality.

(*n* = 60). The regression line with a 95 % confidence interval indicates that there is indeed a positive correlation between polyp mortality and 2-methylnaphthalene exposure. Some data points have similar values, and hence, overlap in the plot. The vertical red dashed lines indicate the 2-methylnaphthalene category thresholds and coincide with the 5 % and 25 % mortality levels (LC5 and LC25, respectively). The corresponding 2-methylnaphthalene concentrations are 1.25 mg L^*−*1^ and 2.3 mg L^*−*1^, respectively.

The variation in polyp mortality was relatively high with a polyp mortality ranging from 0 % to 50 % for concentrations from 0.9 mg L^*−*1^ to 2.3 mg L^*−*1^, and a polyp mortality ranging from 0 % to 100 % for concentrations between 2.3 mg L^*−*1^ to2.7 mg L^*−*1^. The mortality ranged from 0 % to 17 % for concentrations of methylnaphthalene ranging from 0.0 mg L^*−*1^ to 0.9 mg L^*−*1^. The confidence bounds correspond to a 95 % confidence interval, computed by bootstrapping (*N* = 10^4^). The best fit curve is a generalized least squares model with a logit link function, yielding *R*^2^ = 0.54 and was constrained to pass through the origin. The dependent variable was binarized before fitting the model, as polyp mortality can be seen as an effect on the organism.

Based on the analysis shown in Fig 5, categories of exposure have been chosen with limits corresponding to LC5 and LC25:

- Non-exposed and lowest exposed corals: *C* ≤ 1.25 mg L^*−*1^
- Intermediate exposed corals: 1.25 mg L^*−*1^ *Ã C* ≤ 2.30 mg L^*−*1^
- Highest exposed corals: *C >* 2.30 mg L^*−*1^

What we here refer to as low, intermediate and high doses, is based on the doses used in the toxicity test to ensure mortality was obtained, and does not refer to what we consider as low, medium, and high concentrations in situ and associated with, e.g., oil spills.

As such, the doses of 2-methylnaphthalene chosen in this study were not primarily chosen to be environmentally relevant. Compared to measured doses of naphthalenes in sea water samples during oil spills, e.g. Deepwater Horizon, the doses used in the present study are considerably higher [40]. Water samples collected and analyzed for oil compounds, including naphthalenes, during or shortly after an oil spill, are generally below the lowest levels of exposure reported in this study. In a study by Guyormach et al. [41], dissolved PAH concentration was monitored during a field trial in the North Sea, with 2 slicks of respectively 1 m^3^ and 3 m^3^ of Grane crude oil. The maximum concentration, for the sum of 1-methylnaphthalene and 2-methylnaphthalene, was below 2 µg L^*−*1^.

### PLS model

For the PLS model, a total of 10 latent variables were retained; cross-validation suggested an essentially flat plateau of model performance for 5 to 20 latent variables. The first six **X** loading vectors are shown in Fig 6 and Fig 7, for white and orange corals, respectively. For reference, some example spectra (i.e., rows in the **X** matrix) taken from unexposed and high exposure coral samples are shown in S2 Fig. The figures indicate which parts of the spectrum is important for explaining covariance between spectra and 2-methylnaphthalene concentration and polyp mortality. Several spectral features, i.e., peaks and dips, can be observed across the spectrum, especially for higher order loadings. Loading 5 and 6 exhibit high frequency noise, indicating that one is approaching an appropriate cutoff in number of latent variables. Some features appear to be present in both color morphs (e.g., the peak/dip at 650 nm for loading 5 in both color morphs), whereas others are observed only in one color morph (e.g., for white corals, a dip is observed in loading 3 at 540 nm, but no such dip is observed for orange corals). Each spectral dip or peak (note that the sign of loading vectors is arbitrary) can potentially be interpreted as absorption by a chemical compound in the coral. It is of interest to examine further the sources of these spectral features, as it would enable the creation of more robust interpretation models for hyperspectral imagery.

**Fig 6.** PLS X loadings for white corals.

The PLS **X** loadings show which parts of the spectrum are significant in explaining covariance between spectra and 2-methylnaphthalene concentration and polyp mortality. Higher order loadings carry less significance in explaining the variation in the corals exposure doses and mortality; this is a property of the PLS algorithm.

**Fig 7.** PLS X loadings for orange corals.

Same as Fig 6, but created using data from orange corals. A notable difference is the significant noise in loading 6, indicating that explanatory power is decreasing at this point.

After fitting the PLS model to training data, **X** scores were extracted. The scores correspond to coordinates in the latent variable space. The first three score components for white coral data can be seen in Fig 8. One dot in the scatter plot corresponds to one coral spectrum, i.e., one hyperspectral image pixel. While the color scale in the figures is continuous, only 20 distinct values appear. This is due to the experimental setups, where each of the five treatment groups consisted of four replicates, giving a total of 20 exposure beakers containing 3 coral samples each. Measurements of methylnaphthalene concentration were done for each replicate.

**Fig 8.** PLS scores for white corals.

(a) shows scores for **X** latent variable 1 vs latent variable 2, (b) shows scores for **X** latent variable 1 vs latent variable 3. One dot corresponds to one spectrum (i.e. one hyperspectral image pixel) taken from a coral.

When examining Fig 8, we observe clustering of low exposure values near the origin, whereas spectra from corals exposed to higher values of 2-methylnaphthalene form scattered clusters outside of the central cluster. We attribute this clustering partly to the 2-methylnaphthalene exposure levels being highly clustered. When rotated in a 3-dimensional view, one can more easily see that the spectra of high exposed corals form a “shell” outside of the central cluster consisting mainly of low exposed corals, and do not take the shape of a simple linear structure. With this in mind, we note that a linear discriminant will not be able to separate the spectra from the lowest and highest concentrations, nor will a linear regression algorithm (including PLS regression) satisfactorily predict 2-methylnaphthalene exposure level. However, as we observe good separation between the non-exposed to lowest exposed corals, the intermediately exposed corals and the highest exposed corals, we have performed spectral classification using the nonlinear classification algorithm SVM.

Score plots analogous to Fig 8, but for orange *L. pertusa*, are given in S3 Fig. Finally, identical score plots but with polyp mortality as the color variable are shown for both color morphs in S4 Fig.

### Classification

The quality of the classification of exposure category is assessed using the metrics *precision*, *P*, and *recall*, *R*, which are frequently used in the context of classification. The metrics are defined as

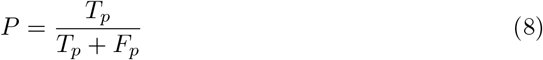

and

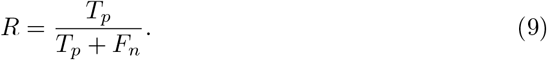

Here, *T_p_* is the fraction of true positive classifications, and *F_p_* and *F_n_* is the fraction of false positive and false negative classification, respectively. Finally, we give the *F* 1-score, which takes both precision and recall into account as the *harmonic* average:

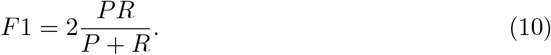

Note that all quantities *T_p_*, *F_n_*, *F_p_*, *P*, *R*, and *F* 1 are defined on the interval [0, 1], and that a high score of *P*, *R*, and *F* 1 equal to 1 is optimal.

The input data is split 80 – 20 into a training data set (80 %) and a test data set (20 %). The split is performed using random stratified sampling, giving each class (no or low, medium, and high exposure) a number of training data samples proportional to the size of the class. The classification model is fit to the training data set, and all classification results shown below are for the test data set.

Results for the three-class case are given in Table 3. The overall picture is that an accuracy in the range 78 % to 97 % is achieved using SVM classification. The poorest performance is found for high exposed corals. This is possibly due to the fact that fewer spectra were collected for these classes (see the “pixels” column of Table 3).

**Table 3.** Summary of classification results for single spectra, using the processing pipeline shown in Fig 4. *P* denotes precision, *R* recall, and *F* 1 is the harmonic average of *P* and *R*.

| Color morph | Exposure | Pixels | *P* | *R* | *F1* |
| --- | --- | --- | --- | --- | --- |
| White | Low | 1252 | 0.87 | 0.95 | 0.91 |
|  | Medium | 457 | 0.87 | 0.80 | 0.83 |
|  | High | 440 | 0.92 | 0.78 | 0.84 |
|  | Total/avg | 2149 | 0.88 | 0.88 | 0.88 |
| Orange | Low | 512 | 0.85 | 0.98 | 0.91 |
|  | Medium | 235 | 0.94 | 0.79 | 0.86 |
|  | High | 216 | 0.97 | 0.78 | 0.86 |
|  | Total/avg | 963 | 0.90 | 0.89 | 0.89 |

Classification results for the three-class case in terms of precision (*P*), recall (*R*), and *F1*-score. An overall *F1* score of 0.88 and 0.89 is achieved for white and orange corals, respectively. The “pixels” column shows the number of spectra collected for each class. Note that average values refer to the *harmonic* average.

#### Per-organism classification

Obviously, each pixel of each coral cannot have its exposure level be classified separately. Hence, following classification of spectra, i.e., hyperspectral image pixels, we classify the corals at the *sample* level. This is done by the majority vote algorithm: the coral sample is assigned to the exposure category that most of its pixels are. In this manner, spatial information is used to improve classification, without resorting to advanced image processing algorithms. When this is done, 100 % of all samples are classified in the correct exposure category.

An illustration of this final classification step is shown in Fig 9. Here, images of two corals for each of the three exposure categories are shown, with classified pixels shown in the row below. A few scattered pixels are classified incorrectly; e.g., in the center image, a few orange and black pixels are interspersed within the red pixels. By using the majority vote algorithm within each connected region, each *Lophelia pertusa* sample is classified correctly. This illustrates the power of spatial information when classifying hyperspectral images. Finally, we note that the spatial pattern below is valuable for environmental monitoring, as this technique can be used directly to generate maps of exposed corals.

**Fig 9.** Classification of organisms.

Two example corals from each exposure category: no-or low exposure (left), medium exposure (center), and high exposure (right). Note that the oddly colored patches in the lower left corner (left image) and upper left corner (center image) are caused by overexposure of a mechanical part holding the corals. The lower row shows classification into no- or low exposure (orange), medium exposure (red), and high exposure (right).

### Reference Alive corals

Finally, the 6 reference alive *L. pertusa* samples (2 orange and 4 white) that were in no way subjected to the handling and experimental conditions in the toxicity test, were scanned with the UHI. Classification on these corals were done with the SVM classifier model trained on data from the main treatment groups C0–C4.

The spectra taken from the orange reference corals classify with a precision of 1.00 and a recall of 0.95, whereas spectra from the white reference corals classify with a precision of 1.00 and a recall of 0.88. This is consistent with the classification results of single spectra in the results above, and gives a verification that the experimental conditions have not systematically modified the spectra, at least with respect to classification results.

Finally, at the organism level, 100 % of reference coral samples were classified in the category of nonexposed-or lowest exposed corals. This is correct, as the reference corals were kept separately from the coral samples exposed to 2-methylnaphthalene, and should thus be representative of healthy corals in the wild.

## Conclusion

Underwater hyperspectral imaging has been shown capable of classifying the cold water coral *L. pertusa* according to their individual exposure to the toxic petroleum compound 2-methylnaphthalene, when categorized according to lethal concentration levels LC5 and LC25 (5% and 25% mortality, respectively). A classification model consisting of projection to latent structures followed by a support vector machine classifier achieves a classification score of 73% to 100% correctness for single spectra. When exploiting the spatial information in the hyperspectral images, a full 100% of *L. pertusa* samples are classified correctly under laboratory conditions. The model has been verified with hyperspectral images of reference coral samples not exposed to any toxic compound.

### Future work

Scientists have often struggled to provide an integrated and non invasive assessment of coral health status. In that respect, this study represents the first step towards a non-invasive, automated method for in situ mapping of deep-water coral condition. In order to develop exposure mapping using underwater hyperspectral imaging into a field ready method, several challenges remain. Such a tool would be more valuable if several indicator species can be used, not just *L. pertusa*. Further studies should be conducted on environmentally relevant doses of relevant mixtures of toxicants, as these are encountered following oil spills. In a coral physiology perspective, future studies should focus on assessing the underlying biochemical mechanisms for the mortality-linked spectral features, analogous to the widely discussed loss of colors (coral bleaching) caused by the death of symbiont chlorophyll containing algae of tropical corals. Determining whether coral mortality induced by other causes than toxicants, e.g., ocean acidification and smothering by drill cuttings, can be measured using the same hyperspectral technique may lend more generality to the method.

**Fig 10.** Example spectra from hyperspectral images of the two color morphs of *Lophelia pertusa*, exposed and unexposed to 2methylnaphthalene. Reflectance spectra of white unexposed corals (top left), orange unexposed corals (top right), white high exposure corals (bottom left), and orange high exposure corals (bottom right). For each subfigure, all three spectra are taken from the same specimen.

**Fig 11.** PLS X scores for spectra from orange corals.

**Fig 12.** PLS X scores for spectra from orange corals.

The color map indicates the polyp mortality.

**Fig 13.** PLS X scores for spectra from white corals.

The color map indicates the polyp mortality.

**Fig 14.** Coral sampling equipment.

ROV with fish net was used. Note that this photo was captured at another field campaign using the same equipment as presented in this study.

**Fig 15.** Experimental setup for UHI image acquisition.

The upper image shows coral samples placed on grids at the bottom of a water filled tank. The UHI with lamps were attached to a scanning mechanism, and was set to image the tank bottom scenery while moving in a “push broom” fashion, keeping constant speed and distance to the tank bottom. A Spectralon reference plate and a PVC plate was used to correct for the inherent optical properties of the water. The lower photo details three coral samples attached to a steel grid.

## Supporting information

S1 Table Measured 2-methylnaphthalene concentration in stock bottles and exposure beakers

Table 4 shows the concentration of 2-methylnaphthalene measured in water samples collected from stock solutions every 24 h during the 96 h exposure period. The average (avg) value and corresponding standard deviation (std dev) are also given. Nominal concentrations, *C*_nom_, are listed for each treatment group. Each water sample was performed in duplicate, with each duplicate labeled I or II in the table.

**Table 4.** Measured 2-methylnaphthalene concentration (mg L^*−*1^) in stock bottles.

| TG<br>$C_{\text{nom}}$<br>Time (h) | C0 | | | C1 | | | C2 | | | C3 | | | C4 | | |
| --- | --- | --- | --- | --- | --- | --- | --- | --- | --- | --- | --- | --- | --- | --- | --- |
|  | I | II | avg | I | II | avg | I | II | avg | I | II | avg | I | II | avg |
| 0 | 0.0 | 0.0 | 0.0 | 1.4 | 1.3 | 1.3 | 3.2 | 3.2 | 3.2 | 3.0 | 3.0 | 3.0 | 7.9 |  | 7.9 |
| 24 | 0.0 | 0.0 | 0.0 | 1.5 | 1.5 | 1.5 | 2.9 | 2.8 | 2.8 | 1.8 | 1.8 | 1.8 | 7.8 | 8.2 | 8.0 |
| 48 | 0.0 | 0.0 | 0.0 | 1.3 | 1.3 | 1.3 | 2.4 | 2.5 | 2.4 | 5.0 | 4.4 | 4.7 | 10.2 | 7.3 | 8.8 |
| 72 | 0.0 | 0.0 | 0.0 | 1.1 | 1.0 | 1.1 | 2.1 | 1.9 | 2.0 | 4.7 | 4.3 | 4.5 | 7.4 | 6.1 | 6.7 |
| avg | 0.0 | 0.0 | 0.0 | 1.3 | 1.3 | 1.3 | 2.6 | 2.6 | 2.6 | 3.6 | 3.4 | 3.5 | 8.3 | 7.2 | 7.9 |
| std dev | 0.0 | 0.0 | 0.0 | 0.2 | 0.2 | 0.2 | 0.5 | 0.6 | 0.5 | 1.5 | 1.3 | 1.4 | 1.3 | 1.1 | 0.8 |
| % of nominal |  |  |  | 125 |  |  | 115 |  |  | 70 |  |  | 98 |  |  |

The bottom three rows indicate average, standard deviation, and percentage of nominal concentration for each column, respectively. The missing value in column C4 II was not analyzed.

Table 5 shows the same results, but for the exposure beakers. The exposure was performed with 3 corals in each exposure beaker, across 4 replicates for each of the 5 treatment groups.

**Table 5.**
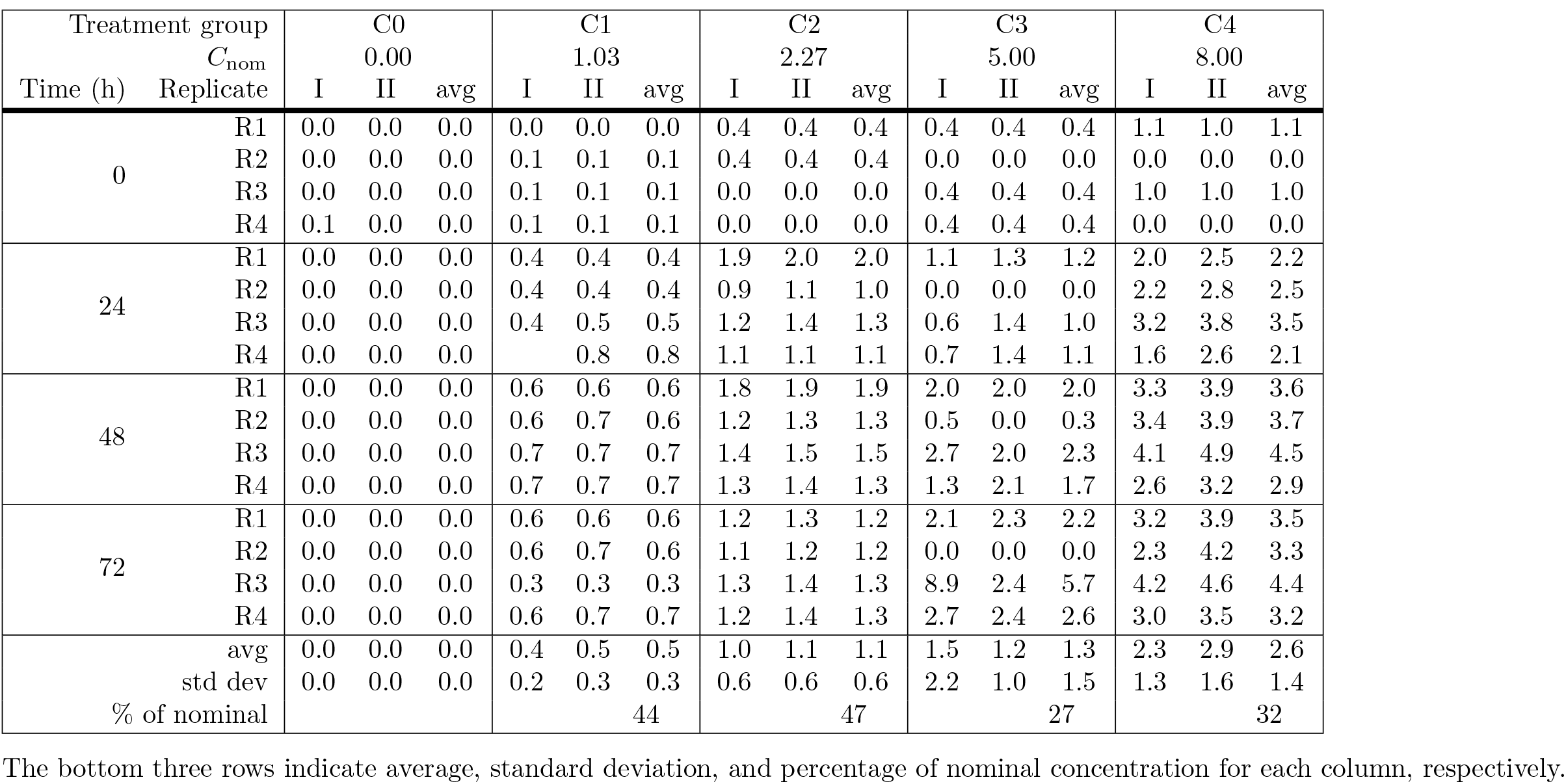
Measured 2-methylnaphthalene concentration (**mg L^*−*1^**) in exposure beakers.

Regarding the analyses performed on the 2-MN water samples, the limit of quantification is 0.1 mg L^*−*1^, and the uncertainty is 20 % at the limit of quantification. This uncertainty decreases above the limit of quantification. Additional uncertainty is expected as the magnetic stirring mechanism cannot operate at high frequency, as this would disturb the coral samples.

S2 Fig Example spectra of corals subject to no or high exposure

Spectra for both orange and white color morphs are given in Fig 10.

S3 Fig PLS X scores for orange corals

PLS **X** scores for orange corals are given in Fig 11. The results correspond exactly to those presented in Fig 8, except that Fig 11 below presents results for the orange color morph.

S4 Fig PLS X scores for corals with mortality coloring

PLS **X** scores colored with the mortality variable for orange and white corals are given in Fig 12 and Fig 13, respectively. The results correspond exactly to those presented in Fig 8 and 11, except for the coloring of the dots.

S5 Fig Equipment used for sampling of corals

The ROV was maneuvered towards the rocky wall growing *L. pertusa* colonies, and the fish net was carefully poked towards the corals from below. This resulted in pieces of the coral colony breaking off and landing in the fish net, as shown in Fig 14. Precaution was taken to avoid any accidental damage of adjacent colonies.

S6 Fig Experimental setup for UHI image acquisition

Fig 15 shows the experimental set-up for hyperspectral imaging. The photo corresponds to schematic sketch given in Fig 3.

## Acknowledgements

This study is part of the joint industry project “New technology and methods for mapping and monitoring of seabed habitats”, financed by the Research Council of Norway (RCN), Statoil, ConocoPhillips Skandinavia, Dea Norge, ENI Norge, Lundin Norway, Total E&P Norge, Norwegian Deepwater Programme, Ecotone, and Akvaplan-niva (RCN project number 235440/E30). The exposure experiment received financial support from the project “Species Sensitivity Distribution for Deep Sea Species and Toxicity of continuous and spiked exposures to crude oil at 1 atm for deep sea species”, led by Akvaplan-niva and financed by the American Petroleum Institute (API). The funders had no role in study design, data collection and analysis, decision to publish, or preparation of the manuscript’ should be used. Ecotone AS, Statoil and Akvaplan-niva AS provided support in the form of salaries for authors PAL, IMH, LMSA, IE, RP, LT, LC and JB but did not have any additional role in the study design, data collection and analysis, decision to publish, or preparation of the manuscript. The specific roles of these authors are articulated in the “author contributions” section.

The authors thank the captain and crew of R/V Gunnerus for collection of coral samples and Johanna Järnegren at the Norwegian Institute for Nature Research (NINA) for advice on coral sampling.

Advice from Geir Johnsen at the Norwegian University of Science and Technology (NTNU) on set-up of the hyperspectral imager rig is highly appreciated.

The personnel at Akvaplan-niva Research and Innovation Station Kraknes (RISK) are acknowledged for facilitating the experiments.

The authors thank Ingvild Andersson at NTNU for valuable discussions on the hyperspectral data results.

